# Tropical invertebrate community assembly processes are robust to a gradient of land use intensity

**DOI:** 10.1101/2023.01.30.526177

**Authors:** Natasha R. Granville, Maxwell V. L. Barclay, Michael J. W. Boyle, Arthur Y. C. Chung, Tom M. Fayle, Huai En Hah, Jane L. Hardwick, Lois Kinneen, Roger L. Kitching, Sarah C. Maunsell, Jeremy A. Miller, Adam C. Sharp, Nigel E. Stork, Leona Wai, Kalsum M. Yusah, Robert M. Ewers

## Abstract

Understanding how community assembly processes drive biodiversity patterns is a central goal of community ecology. While it is generally accepted that ecological communities are assembled by both stochastic and deterministic processes, quantifying their relative importance remains challenging. Few studies have investigated how the relative importance of stochastic and deterministic community assembly processes vary among taxa and along gradients of habitat degradation. Using data on 1,645 arthropod species across seven taxonomic groups in Malaysian Borneo, we quantified the importance of ecological stochasticity and of a suite of community assembly processes across a gradient of logging intensity. The relationship between logging and community assembly varied depending on the specific combination of taxa and stochasticity metric used, but, in general, the processes that govern invertebrate community assembly were remarkably robust to changes in land use intensity.

## Introduction

Community assembly processes drive biodiversity patterns, and a key goal in community ecology is to quantify the relative importance of different community assembly processes. Simultaneously, it is well-documented that land use changes impact biodiversity (Newbold et al. 2015), but it is much less well-known that they can also affect community assembly mechanisms (Wearn et al. 2019). Currently, therefore, we have a strong awareness of the patterns of biodiversity change that are generated by land use change (Newbold et al. 2015), but little understanding of the extent to which the fundamental community assembly processes that create that change are impacted. Moreover, the specific assembly processes generating ecological communities may differ among taxa due to differences in trait evolution (Weiher et al. 2011), meaning studies examining land use impacts on the assembly processes of mammals (e.g.Wearn et al. 2019) may provide little insight into the impacts on other taxa. Attempts to rely on natural ecological processes to restore biodiversity rely, by definition, on naturally occurring community assembly processes (Palmer et al. 1997, Hilderbrand et al. 2005). It is therefore of fundamental importance that we gain a deeper understanding of whether those assembly processes in modified habitats are the same or different to those observed in primary habitats.

Community assembly involves a combination of determinism and stochasticity, there has been long-standing debate over their relative influences (Connor and Simberloff 1979, Chase and Myers 2011). Stochastic assembly generates community diversity patterns indistinguishable from those generated by random chance alone (Hubbell 2001, Chase and Myers 2011, Ning et al. 2019) and can involve random variation around species average demographic rates due to the probabilistic nature of demographic processes like birth, death and migration (Adler et al. 2007, Shoemaker et al. 2020). Deterministic community assembly, on the other hand, involves non-random, niche-based processes such as environmental filtering and biotic interactions (Chase and Myers 2011). Determinism and stochasticity are opposite ends of a continuum, with real-world communities existing somewhere between these extremes (Gravel et al. 2006, Kitching 2013). While previous studies have investigated the balance between stochasticity and determinism in different environments (e.g. Ellwood et al. 2009, Caruso et al. 2012, Shipley et al. 2012, Ortega-Martínez et al. 2020, Valdivia et al. 2021), few have sought to characterise the effect of land use change on this balance.

Therefore, an understanding of the effects of land use change on the relative importance of stochastic and deterministic community assembly processes is largely absent, despite its potential importance. It is often suggested that community assembly processes exhibit hysteresis, meaning that the pathway to reversing damage done to an ecosystem is not the same as the pathway to creating that damage (Beisner et al. 2003). Recovering an ecological community in a modified habitat could rely on different assembly processes than those that exist in primary habitats (Andersen et al. 2009, Suding and Hobbs 2009). Therefore, our understanding of primary community assembly may give information that at best is irrelevant, or at worst directly misleading, when it comes to planning the restoration of modified communities. We therefore need to characterise how community assembly mechanisms might change across gradients of habitat degradation.

Logging is a major driver of habitat degradation across many of the world’s most productive and biodiverse tropical forest ecosystems (Laurance 2015). The tropical forests of Borneo have been subject to rapid and widespread logging since the early 1970s. Between 1973 and 2010, there was an estimated 30 % decline in the extent of Borneo’s intact forests (Gaveau et al. 2014). Logging often results in a heterogeneous landscape, with habitat patches connected spatially but affected by different logging intensities (Berry et al. 2008). This can result in gradients of disturbance intensity, which are a frequent consequence of land use change in the tropics (Wearn et al. 2019).

There is uncertainty over how logging might affect stochastic and deterministic community assembly processes, with little in the way of direct evidence. Kitching et al. (2013) showed a distance-turnover relationship (decreasing community similarity with increasing geographic distance) for moth communities in primary forest, but such patterns were largely absent in logged forests, suggesting that stochastic turnover in plant species can drive deterministic changes in the niche dimensions available for moths in primary forests, but not in logged forests. Döbert et al. (2017) showed that understorey plant communities in tropical Bornean forests tend to be more stochastically assembled at higher logging intensities, whereas Wearn et al. (2019) showed that deterministic environmental control on community assembly is higher in logged forest compared to old-growth forest. There is broad conceptual and empirical support for deterministic environmental filtering on community assembly becoming more important in harsher environmental conditions (Chase 2007, Lepori and Malmqvist 2009, Chase and Myers 2011, Ding et al. 2012, Wearn et al. 2019, Li et al. 2021, Hu et al. 2022). We therefore hypothesise that stochasticity will become less important with increasing logging intensity.

There is further uncertainty over how taxonomic groups might differ in their community assembly processes. There is broad support for vastly different taxonomic groups showing variation in the balance between stochasticity and determinism. For example, Powell et al. (2015) found substantial differences in the assembly of soil bacterial and fungal communities, due to differences in dispersal capacity between these groups. Trophic position is also likely to play a role in mediating these differences between taxonomic groups; Keppeler et al. (2016) indicated differences in assembly processes between foraging bird and fish communities. Among comparatively more similar taxa, Thompson and Townsend (2006) showed that trophic groupings and species traits determine the relative importance of stochasticity and determinism in macroinvertebrate communities. Generalist taxa have been shown to contribute more to stochastic processes, and specialist taxa to contribute more to deterministic processes in microbial communities, because specialists tend to have narrower tolerance to environmental changes (Liao et al. 2016, Xu et al. 2022). Similarly, studies on microbial communities have also shown that rare taxa, which have narrower niche breadths, are more deterministically assembled while abundant taxa with wider niche breadths are more stochastically assembled (Gao et al. 2020, Yang et al. 2022). There has not, to our knowledge, been a similar study directly quantifying among-taxa variation in stochastic and deterministic assembly processes for invertebrate communities.

Stochasticity indices have been widely used in studies on microbial communities to quantify the importance of ecological stochasticity (e.g. Jiao et al. 2020, Le Moigne et al. 2020, Sun et al. 2021, Trego et al. 2021, Zhou et al. 2022, Wang et al. 2022). These indices compare observed community dissimilarity to the neutral expectation. Neutral models generally assume that all individuals within a feeding guild have equal chances of birth, death and migration, regardless of their taxonomic identity (Hubbell 2001). Significant deviations from neutral models of random community assembly can indicate that other processes, such as selection, are involved in structuring the community (Burns et al. 2016). The relative importance of classes of community assembly processes can be inferred from patterns of taxonomic and phylogenetic diversity (Ning et al. 2020). Phylogenetic and taxonomic turnover can indicate whether assembly processes are driving communities to be more heterogeneous (high turnover) or homogeneous (low turnover), and comparing these patterns with neutral expectation can indicate the relative importance of stochasticity (Ning et al. 2020).

There is a long-standing need to evaluate the relative importance of stochastic and deterministic processes along environmental gradients and among taxa (Weiher et al. 2011). Here, we address that knowledge gap by using a variety of indices to quantify the relative contribution of stochasticity to community assembly for seven invertebrate taxa across a gradient of logging intensity. We also quantify the relative importance of a suite of community assembly processes across the logging gradient. Our data encompass a comprehensive gradient of logging intensity, from areas that have never experienced logging to areas that have been salvage logged. We quantified community assembly for a range of invertebrate taxa including three groups of Coleoptera (beetles), along with Formicidae (ants), Lepidoptera (moths), Orthoptera and Araneae (spiders). Together, these taxonomic groups encompass a range of feeding guilds and are of immense ecological importance (Barlow and Woiwod 1989, Didham et al. 1998, Grimaldi et al. 2005, Nyffeler and Birkhofer 2017, Oumarou Ngoute et al. 2020). We use our data to test two hypotheses: (1) stochastic assembly will decrease in importance as logging intensity increases, as logged forest should have stronger environmental filtering; and (2) stochastic assembly will have a lower relative importance for trophic specialists, compared to trophic generalists, because trophic specialists are expected to be more strongly assembled by selective environmental filtering. Finally, we investigate whether the relative importance of a suite of different community assembly processes varies across a gradient of logging intensity for different invertebrate taxa.

## Methods

### Data collection

The study sites were located within the Stability of Altered Forest Ecosystems (SAFE) project (4° 38′ N to 4° 46′ N, 116° 57′ to 117° 42′ E), a large-scale ecological experiment encompassing a gradient of land use intensities in the lowland tropical forests of Sabah, Malaysian Borneo (Ewers et al. 2011). We used data from 14 out of the 17 experimental sampling blocks at SAFE (Ewers et al. 2011), excluding three blocks located in oil palm plantation. Ten sampling blocks were located in twice-logged forests and four were located in protected areas. Two of these protected area blocks were in the Maliau Basin Conservation area and have never experienced logging, while the other two had experienced light logging through both legal and illegal processes. Each sampling block comprised a set of 4 – 43 sampling sites (mean = 19) and covered a spatial area of 4 – 229 ha (mean = 56). We grouped invertebrate samples collected within each block which we considered as one local community for analysis. The aggregation of all local communities across all sampling blocks was considered to represent the metacommunity.

Above-ground carbon density (ACD), calculated from LiDAR surveys and summarized at 1 ha resolution in 2014, was used to quantify logging intensity (Jucker et al. 2018, Swinfield et al. 2020): a higher above-ground carbon density corresponds to a lower logging intensity. ACD was log-transformed to generate a more uniform spread of logging intensity values. The sampling blocks covered a wide range of logging intensities, with average above-ground carbon densities ranging from around 15 t.C.ha^-1^ in heavily logged locations, to over 200 t.C.ha^-1^ in the protected areas. Not all invertebrate taxa were sampled at the same subset of sampling points per block. For analysis, then, we calculated the average ACD per sampling block separately for each taxon, taking as inputs the ACD values for the specific subset of sampling sites where that taxon was collected.

We combined community composition data collected from seven invertebrate taxa: three groups of beetles, plus ants, moths, spiders and Orthoptera. Different groups had different sample sizes and not all groups were sampled in all 14 sampling blocks (Table S1, Figure S1). Beetles were sampled between 2011 and 2013 using combination pitfall-malaise traps in all 14 sampling blocks. Three different groups of beetles were sampled: Curculionoidea (weevils), Staphylinidae (rove beetles) and Scarabaeoidea (scarabs) (Sharp et al. 2018, 2019). Because of differences in their feeding guilds, each group was considered a separate taxon and was analysed separately: weevils are predominantly herbivorous; most scarabs in our dataset are dung-feeders; and rove beetles can belong to several feeding guilds. Beetles were identified primarily to morphospecies, except some scarabs which were identified to species. Ants were sampled between December 2011 and June 2012 in 12 sampling blocks using 12cm x 14cm plastic cards, which were laid flat in the leaf litter and baited with 30 compressed dried earthworm pellets. The number of ants entering each card was observed and recorded for 40 minutes, and individuals were identified to morphospecies (Fayle et al. 2019). Moths were sampled in 2014 using UV light traps which were run overnight in 8 sampling blocks. Moths were identified where possible and separated into morphospecies using morphology (Maunsell et al. 2020b). Spider abundance data were collected in 2015 in 10 sampling blocks by beating plant foliage for 20 minutes at each site. Spiders were identified to family, then separated into morphospecies by genitalia dissections and DNA barcoding using the CO1 gene (Maunsell et al. 2020a). Finally, Orthoptera were sampled in 2015 by sweep netting along 100 m transects in 6 sampling blocks. Orthoptera were identified to family, then separated into morphospecies using identification guides (Hardwick et al. 2022).

### Stochasticity metrics

There are different ways for communities to express stochasticity, so there is value in assessing multiple metrics of stochasticity on the same communities (Vellend and Agrawal 2010). We quantified stochasticity using three null-model based mathematical frameworks that summarise stochasticity both at the community level and at the level of individual species. All three stochasticity metrics were calculated from a separate community composition (site × species) matrix for each of the seven taxa. We calculated stochasticity metrics for all sites in the composition matrix, and grouped sites together by sampling block to calculate the mean of each stochasticity metric for each sampling block.

First, we employed the normalised stochasticity ratio (NST) (Ning et al. 2019) to assess the relative importance of ecological stochasticity. NST values are normalised on a scale from 0 to 1, with 0.5 as the boundary between more stochastic (>0.5) and more deterministic (<0.5) community assembly. NST is based on pairwise community dissimilarity measures, for which there are many competing metrics. We used Ružička dissimilarity which was shown to have the highest accuracy and precision when the NST was developed (Ning et al. 2019). NST compares the observed dissimilarity of the real community with the null expected dissimilarity for 1,000 randomised communities (Ning et al. 2019). To generate the random metacommunities for the null expectation, the total number of species was fixed as observed, and within each local community, individuals were drawn at random from the metacommunity with probabilities proportional to their regional occurrence frequencies.

Second, to test the effect of different assumptions in the null modelling framework, we also used the modified stochasticity ratio (MST) which transforms NST under the assumption that observed community similarity is equal to the mean of the null expected community similarity under stochastic assembly. MST is also normalised from 0 to 1 with values closer to 1 indicating a higher importance of ecological stochasticity. The NST and MST metrics assess ecological stochasticity at the community level, whereas our third metric, the neutral taxa percentage (NTP) assesses stochasticity at the level of individual species (Burns et al. 2016). NTP is the proportion of species with occurrence frequencies that could be predicted by Sloan’s neutral model (Sloan et al. 2006, 2007). This model assumes that assembly is driven solely by chance and dispersal. It predicts the relationship between the occurrence frequencies of taxa in a set of local communities and their abundances across the metacommunity (Sloan et al. 2006, 2007, Burns et al. 2016). The model is fitted to the frequency and abundance of species in a metacommunity by a single parameter which describes the migration rate (the probability that loss of an individual in a local community will be replaced by dispersal from the metacommunity rather than by reproduction within the local community) (Sloan et al. 2006, 2007, Burns et al. 2016). ‘Neutral taxa’ are those whose observed occurrence frequencies are within one confidence interval of that which would be expected by Sloan’s neutral model (Burns et al. 2016). The proportion of neutral taxa was weighted according to the abundance of individuals in each taxon, and was used to assess the relative importance of ecological stochasticity at the level of individual species (Burns et al. 2016).

### Community assembly processes

To further characterise the drivers of community assembly across the logging gradient, we quantified the relative importance of a suite of assembly processes: dispersal limitation, homogenising dispersal, drift, homogeneous selection and heterogeneous selection (Stegen et al. 2013, Ning et al. 2020). Heterogeneous selection is selection under heterogeneous abiotic and biotic conditions that leads to high phylogenetic compositional variation among local communities. Homogeneous selection takes place under homogeneous conditions and leads to low phylogenetic compositional variation. Dispersal limitation refers to the situation where low levels of dispersal among local communities constrains the exchange of organisms between these communities, leading to high levels of spatial turnover. Homogenising dispersal is the opposite situation where high levels of dispersal result in little turnover among communities. Communities that are not dominated by selection or dispersal are designated as being governed by drift (Stegen et al. 2013, Ning et al. 2020).

The proportional contribution of each of these five community assembly mechanisms was inferred through phylogenetic bin-based null model analysis (iCAMP) (Stegen et al. 2013, Ning et al. 2020). For each of the seven groups of taxa, we used iCAMP to calculate the relative importance of each process for all pairwise comparisons between all sites. We then grouped sites by sampling block and calculated the mean relative importance of each process for each taxa x sampling block combination.

The iCAMP framework divides taxa into groups (‘bins’) based on their phylogenetic relationships, then identifies the dominant community assembly process in each bin. Phylogenies were not available for the taxa included in this study, so we used taxonomy trees as proxies for phylogenies. Taxa were divided into bins based on taxonomic identity (Table S2, Figure S3), and bins with fewer than the minimum number of taxa (n = 9) were merged into the bin to which they were most closely related (Ning et al. 2020). The number of randomisations used for the null model analysis was 500.

To calculate the relative importance of community assembly processes, iCAMP uses null model analysis of phylogenetic diversity (beta net relatedness index, βNRI) and taxonomic β-diversity (modified Raup-Crick metric, RC) to identify the process governing each bin. According to the framework developed by Ning et al. (2020); for each bin, the fraction of pairwise comparisons between sites with βNRI < −1.96 was considered as the percentage contribution of pairwise comparisons dominated by homogeneous selection, and those with βNRI > +1.96 as the percentage contribution of heterogeneous selection. Pairwise comparisons with −1.96 ≤ βNRI ≤ +1.96 were further partitioned using RC. The fraction of pairwise comparisons with RC < −0.95 indicates the percentage contribution of homogenising dispersal, while those with RC > +0.95 indicates the percentage contribution of dispersal limitation. The remaining pairwise comparisons with βNRI ≤ |1.96| and RC ≤ |0.95| were considered to represent the contribution of drift, which includes ecological drift and other processes such as diversification, weak selection and weak dispersal. The fractions of each process across all bins were then weighted by the relative abundance of each bin (Stegen et al. 2013, Ning et al. 2020).

### Statistical analysis

All data analysis was conducted using R version 4.1.0 (R Core Team 2022). NST and MST stochasticity metrics were calculated using the *NST* package (Ning et al. 2019), while NTP and iCAMP metrics were calculated using the *iCAMP* package (Ning et al. 2020).

To see how species identity changed across the gradient of logging intensity, we compared the Bray-Curtis similarity of each sampling block to old-growth forest using the *vegan* package (Oksanen et al. 2022), and used beta regression (Cribari-Neto and Zeileis 2010) to assess the effect of ACD on Bray-Curtis similarity. We also analysed species richness trends among taxa and across logging gradients, and compared these trends to those of the stochasticity metrics to see whether the metrics were affected by species richness.

We calculated the mean of each stochasticity metric (NST, MST and NTP) and of each community assembly process (heterogeneous selection, homogeneous selection, dispersal limitation, homogenising dispersal and drift) for each taxa x sampling block combination. We estimated the 95 % quantiles of these means using bootstrapping; we sampled the values used to calculate the mean 1000 times with replacement, then took the 5^th^ and 95^th^ quantiles of this distribution.

To compare the relative importance of stochasticity among taxa we used one-sample *t*-tests to compare the overall unweighted mean NST, MST and NTP across all taxa to 0.5, which we used as a boundary point separating stochastic (>0.5) from deterministic (<0.5) community assembly. We also used ANOVA to test for differences in NST, MST and NTP among taxa. To test the hypothesis that trophic generalists (ants and rove beetles) would be more stochastic than trophic specialists (moths, Orthoptera, spiders, scarabs and weevils), we used beta regression with a single categorical predictor that describes whether the taxon is considered a trophic generalist or specialist. We fitted three separate beta regression models, each with a stochasticity metric as the response variable. We broadly categorised each taxonomic group into trophic generalists or specialists based on trophic level. Ants were considered generalists because their diets can range from almost herbivorous to omnivorous and fully predatory (Blüthgen et al. 2004). Many rove beetles are generalist predators, though some can belong to other feeding guilds (Méndez-Rojas et al. 2021), so we also classified rove beetles as trophic generalists. The Orthoptera and weevils included in our data were mainly herbivorous (Sharp et al. 2019, Hardwick et al. 2022), so we classified them as trophic specialists. Spiders were all predatory (Russell-Smith and Stork 1995), scarabs were primarily coprophagous (Sharp et al. 2018) and moths were herbivorous as larvae and nectarivorous as adults (Romeis et al. 2005), so we classified these groups as trophic specialists for our analysis.

To investigate the effect of logging on the importance of each stochasticity metric we used beta regression, with the mean ACD of each block as the predictor. To see how this relationship varies among taxa, we fitted another beta regression model for each stochasticity metric with taxa, ACD and their interaction as predictors of stochasticity. A similar analysis was conducted to investigate the effect of logging on the relative importance of each community assembly process, but Dirichlet regression was used instead of beta regression as Dirichlet regression is appropriate for proportions with more than two categories (Douma and Weedon 2019).

To gain a taxa-independent metric reflecting the overall change in the relative importance of the five community assembly processes across the logging gradient, we combined the slopes for all taxa to give a weighted summary mean of the slopes for each process, using a method similar to a fixed effects meta-analysis, adapted from Borenstein et al. (2010). We also combined the mean relative importance of each of the five processes and each of the three stochasticity metrics to give a weighted summary mean for all taxa using the following method adapted from Borenstein et al. (2010):

For each group of taxa (𝑖), the weight (𝑤) is the inverse of the variance for that group 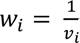 . The variance (𝑣) is the range of the 95 % confidence intervals. 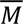 is the overall weighted mean for all groups of taxa combined, it is analogous to the combined effect size in a meta-analysis. 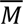 is calculated as 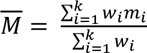 where 𝑚 is the mean relative importance or slope for each group of taxa, analogous to the effect size of each study in a meta-analysis, and 𝑤_𝑖_ is the weight assigned to group 𝑖, and 𝑘 is the number of groups of taxa (𝑘 = 7).

## Results

Across all datasets, our analysis included 32,294 individuals belonging to 1,645 species or morphospecies (Table S1). In general, community similarity between sampling blocks was low. Each of the three beetle taxa, as well as moths, showed a decrease in community similarity compared to old growth forest as logging intensity increased, whereas the remaining three taxa (ants, Orthoptera and spiders) did not (Figure S2).

### Stochasticity metrics

The overall average NST across all taxa was 0.49 (95 % CI = 0.44 – 0.54), which was not significantly different from 0.5 (one-sample *t*-test, *t* = −0.30, df = 76, *p* = 0.76). However, the MST metric indicated that determinism plays a greater role in structuring invertebrate communities than ecological stochasticity (MST: mean = 0.28; 95 % CI = 0.22 – 0.33; *t* = −8.19, df = 76, *p* < 0.0001). In contrast, NTP indicated a greater role of stochasticity (NTP: mean = 0.68; 95 % CI = 0.64 – 0.71; *t* = 10.1, df = 63, *p* < 0.0001). When the overall average NST was weighted by the variance of each taxon, it reduced to 0.39 (95 % CI = 0.07 – 0.83) and MST reduced to 0.10 (95 % CI = 0.01 – 0.75). The unweighted average NTP was not significantly different to the weighted average NTP (0.69, 95 % CI = 0.43 – 0.89) (Figure 1).

**Figure 1.**
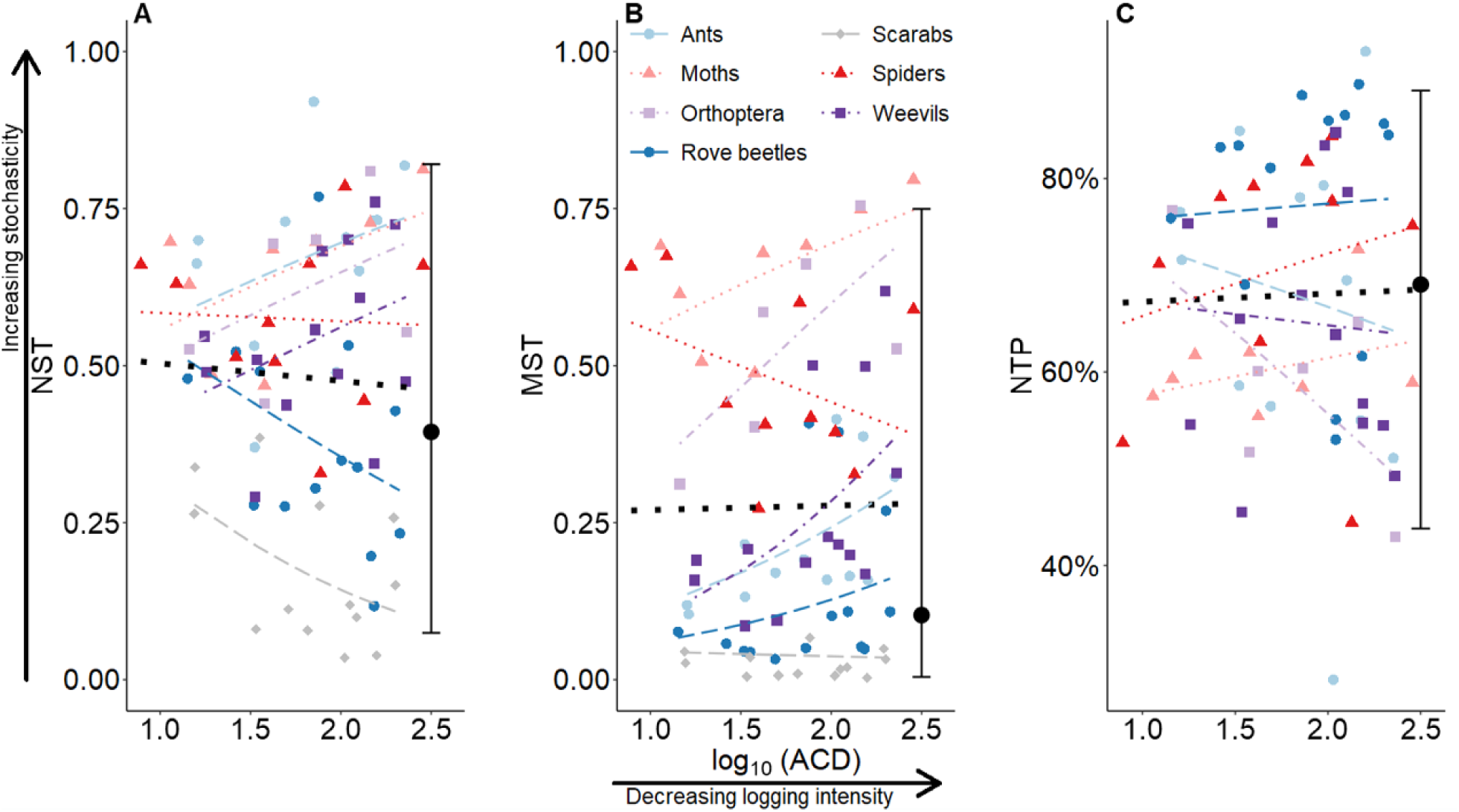
Variation in three different metrics of ecological stochasticity across a gradient of logging intensity for seven invertebrate taxa. Logging is quantified as log_10_-transformed above-ground carbon density (ACD), where a higher ACD corresponds to a lower logging intensity. The thick black dotted regression lines indicate that there was no significant effect of logging on stochasticity when all taxa were combined. The black circles show the weighted average stochasticity metrics for all taxa combined, and the error bars represent 95 % quantiles of these averages. The three stochasticity metrics are: (A) Normalised stochasticity ratio (Ning et al. 2019); (B) Modified stochasticity ratio (Ning et al. 2019); and (C) Proportion of taxa with observed occurrence frequencies predicted by Sloan’s neutral model (Sloan et al. 2006, 2007, Burns et al. 2016).

There were significant differences in the role of stochasticity among taxa (ANOVA, NST: *F*_70, 6_ = 19.7, *p* <0.0001; MST: *F*_70, 6_ = 33.6, *p* < 0.0001; NTP: *F*_58, 5_ = 2.6, *p* = 0.04). On average, ants had the highest NST, whereas scarabs had the lowest NST. Scarabs also had the lowest MST, while moths had the highest MST. Rove beetles had the highest NTP and Orthoptera had the lowest NTP. Trophic generalists had higher NTP than trophic specialists (beta regression, coefficient = −0.39, *z* = −2.56, *p* = 0.02), however there was no significant difference in NST or MST between trophic generalists and trophic specialists (NST: coefficient = −0.22, *z* = −1.02, *p* = 0.31. MST: coefficient = 0.36, *z* = 1.47, *p* = 0.14).

The relative importance of stochasticity was calculated within each sampling block then compared among blocks. Each block had a different level of logging intensity. When all taxa were combined, there was no significant effect of logging on stochasticity (Figure 1. NST: slope = −0.11, *z* = −0.44, *p* = 0.66. MST: slope = 0.03, *z* = 0.12, *p* = 0.91. NTP: slope = 0.04, *z* = 0.20, *p* =0.84). The relationship between logging and stochasticity varied among taxa (Figure 1). But, in general, the slopes for each taxon were not significant, with the only significant relationship being an increase in MST with decreasing logging intensity for ants (slope = 0.89, 95 % CIs = 0.02 – 1.76). Sloan’s neutral model could not be fitted to the scarabs dataset, so scarabs were excluded from the NTP analysis.

### Community assembly processes

Dispersal limitation was the dominant community assembly process when all taxa were combined and weighted by sample size (overall relative importance = 64 %, 95 % CI = 26 – 97). Dispersal limitation was the dominant process for four out of the seven taxonomic groups (rove beetles, scarabs, weevils and ants), with a relative importance ranging from 94 to 97 % (95 % quantiles = 88 - 99). Drift was the dominant process for spiders, Orthoptera and moths; 89 % of spider community assembly (95 % quantiles = 78 - 96) was estimated to be underpinned by drift, while this was estimated to be 75 % for Orthoptera (95 % quantiles = 55 - 87) and 67 % for moths (95 % quantiles = 54 - 78) (Figure 2A; Table S4).

**Figure 2.**
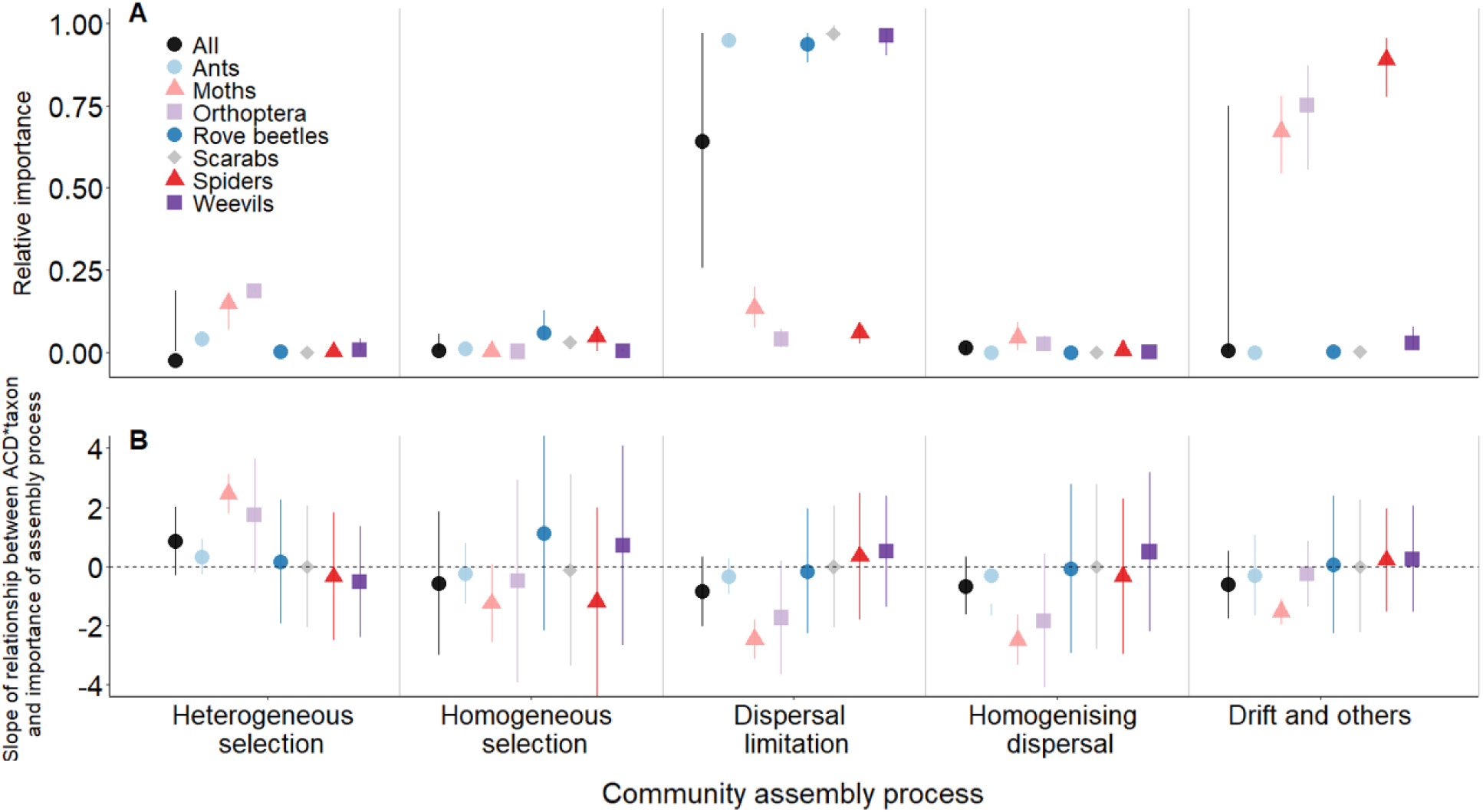
**(A)** Mean relative importance of five community assembly processes for generating the community structure of seven invertebrate taxa, and weighted averages for all taxa combined. Error bars show bootstrapped 95 % quantiles. **(B)** The estimates for the slopes of the relationships between the relative importance of five community assembly processes and above-ground carbon density (ACD). Values are shown separately for each of seven invertebrate taxa, and the black circles show weighted averages for all groups of taxa combined. Positive slopes indicate processes that become less important as logging intensity increases.

When comparing the relative importance of community assembly processes among sampling blocks, each of which had a different level of logging intensity, the relative importance of the five community assembly processes was not significantly affected by logging intensity for most taxa (Figure 2B, Table S5). When all groups of taxa were combined, the weighted mean slopes for each process were not significantly different from zero. The strongest effects were a decrease in the importance of dispersal limitation as logging intensity decreases (−0.85, 95 % CI = −2.01 – 0.32) and an increase in heterogeneous selection with decreasing logging intensity (0.85, 95 % CI = −0.32 – 2.01), but these were not significantly different from zero. The mean effects for homogeneous selection (−0.55), homogenising dispersal (−0.65) and drift (−0.61) were even weaker. The only significant slopes for individual groups of taxa were for moths, which showed a decrease in the importance of dispersal limitation, homogenising dispersal and drift with decreasing logging intensity (Dispersal limitation: slope = −2.45, 95 % CI = −3.11 – −1 .79. Homogenising dispersal: −2.49, 95 % CI = − 3.34 – −1.64. Drift: −1.53, 95 % CI = −1.97 – −1.08) (Figure 2B; Table S5), and an increase in the importance of heterogeneous selection with decreasing logging intensity (Heterogeneous selection: 2.45, 95 % CI = 1.79 – 3.11).

## Discussion

While the relative importance of stochasticity varied among taxa and metrics, in general, the balance between stochasticity and determinism appeared robust to a gradient of land use intensity. At a finer resolution, dispersal limitation, inferred from community patterns, was the dominant assembly process overall. Two out of the three stochasticity metrics (NST and MST) were not significantly different between trophic generalists and trophic specialists. Only the NTP metric supported our hypothesis that trophic generalists would be more stochastic than trophic specialists. In general, for all taxa except moths, land use change had little impact on the relative importance of a suite of community assembly processes. Together, this suggests that, while logging has profoundly negative impacts on biodiversity, it tends to have little impact on the main community assembly drivers for most invertebrate taxa studied.

Overall, there was at best a very weak effect of land use intensity on the role of ecological stochasticity in structuring insect communities. The direction of the relationship between logging and stochasticity varied among taxa, but these relationships were not statistically significant (Figure 1). The relative importance of different community assembly mechanisms was also not significantly affected by logging intensity for six out of seven taxonomic groups (Figure 2B). Moths, however, were the exception, showing changes in community assembly processes (Figure 2B) and turnover in species composition (Figure S2) with logging. For the six taxa that showed no change in community assembly with logging, we might expect to find little evidence of a change in the assembly processes governing invertebrate communities if those communities do not exhibit turnover in species composition across the logging gradient. For three of the taxa (ants, Orthoptera and spiders), this assumption holds true: we found no evidence of changing taxonomic identities across the logging gradient (Figure S2) which aligns well with a lack of change in the assembly processes governing those taxa (Figure 2B). However, all three beetle taxa (rove beetles, scarabs and weevils) did exhibit significant turnover in taxonomic identity as logging intensity increased (Figure S2), which is consistent with previous studies (Hamer et al. 2003, Cleary et al. 2007, Sharp et al. 2019). Yet these taxa did not exhibit significant changes in community assembly metrics across the logging gradient, suggesting that the species turnover was generated by the same ecological processes, regardless of logging intensity. This leads to the general conclusion that, regardless of whether taxonomic identities change, community assembly processes remain robust to changes in above-ground carbon density.

Since the community assembly metrics used here are based on taxonomic and phylogenetic diversity, they could be influenced by changes in species richness, which can change systematically among taxa and along logging gradients (Burivalova et al. 2014). However, we analysed changes in species richness and showed that they are unlikely to underpin our results, as patterns of species richness (Figures S3, S4) differed from those of stochasticity metrics and community assembly processes (Figures 1, 2). This suggests that the trends in stochasticity and community assembly are independent of any trends in species richness.

One possible explanation for why community assembly processes appear to be strongly conserved across the gradient of logging intensity is that our data were collected after logging had taken place. While our study landscape encompasses a very wide range of historic logging intensity, the time delay between the logging event itself and our description of the invertebrate communities means any transitory impacts of logging on community assembly processes would not have been detectable. Mahayani et al. (2020) showed that phylogenetic diversity and community structure of tree communities had recovered 10 years after a single logging cycle in Bornean tropical forest. Therefore, it could be possible that the ecological communities we sampled might have recovered their basic, pre-logging structures so that the community assembly mechanisms in logged forests now resemble those of unlogged forests.

In general, the processes that govern community assembly were robust to logging in our study. However, some relationships were significant for certain combinations of taxa and process. Interestingly, moths showed a decrease in the importance of heterogeneous selection, and an increase in the importance of homogeneous selection and homogenising dispersal, with increasing logging intensity. Together, these trends suggest that logging could be driving the biotic homogenisation of moth assemblages.

When inferring assembly processes from community patterns, dispersal limitation was the most important driver of community assembly for ants and beetles, whereas drift was the main driver of assembly for spiders, Orthoptera and moths (Figure 2A). Spiders, Orthoptera and moths were sampled at fewer sites than ants and beetles (Table S1), so we cannot definitively rule out the possibility that this result may represent a sampling effect. We do note, however, that the scarab community had a lower total number of individuals and a higher number of species than the Orthoptera, suggesting that an undersampling-driven effect should have exerted a greater impact on them than on the Orthoptera. Further, when we conducted a sensitivity analysis by grouping sites within the same block, drift was still the dominant community assembly process for spiders, moths and Orthoptera (Text S1).

The overall importance of dispersal limitation as the dominant community assembly process, especially for ants and beetles highlights the importance of maintaining and, if necessary, restoring landscape connectivity in logged forests (Wearn et al. 2019). We emphasise that dispersal limitation was not directly measured in this study but was inferred from community patterns. Dispersal limitation can result in high spatial turnover in community composition due to low levels of exchange of organisms among local communities (Stegen et al. 2013). The slight increase in dispersal limitation we detected as logging intensity increased, though not significant, is in a direction that is consistent with previous studies suggesting that animal communities in logged tropical forests may experience lower levels of dispersal compared to primary forests (Stratford and Robinson 2005, Laurance et al. 2009, Edwards et al. 2014).

This study has quantified community assembly mechanisms across a gradient of logging intensity for seven groups of invertebrate taxa in Bornean rainforests. The effect of logging on stochasticity, and on different community assembly processes, varied among the different taxa and different metrics of stochasticity, but painted a general picture in which the dominant community assembly mechanisms are not impacted by logging disturbance. Although logging did not alter the balance between stochastic and deterministic community assembly processes for most taxa, we emphasise that logging, and in particular severe logging, profoundly reduces species richness and changes community composition (Thorn et al. 2018). The robustness of invertebrate communities to logging disturbance in our study suggests that knowledge of primary community assembly can be useful in planning the restoration of modified communities as, for six out of seven taxonomic groups, there were not significant changes in assembly processes despite changes in land use intensity.

## Acknowledgements

Tom Swinfield, David Milodowski, Tommaso Jucker, Michele Dalponte and David Coomes generated the above-ground carbon density layer from LiDAR data. TMF was supported by a Czech Science Foundation Standard Grant (19-14620S). Field data collection was funded by Australian Research Council (ARC) Discovery Project Grants (DP140101541 and DP160102078).

## Data accessibility statement

The data that support the findings of this study are openly available on Zenodo at:

https://doi.org/10.5281/zenodo.1323504 (beetles)

https://doi.org/10.5281/zenodo.7011354 (Orthoptera)

https://doi.org/10.5281/zenodo.4247169 (moths)

https://doi.org/10.5281/zenodo.3876227 (ants)

https://doi.org/10.5281/zenodo.4020697 (above-ground carbon density)

The R scripts used for all analyses in this study are openly available on GitHub at: https://github.com/natashagranville/invertebrate-community-assembly

A pre-print of this manuscript was uploaded to BioRxiv:

Natasha R. Granville, Maxwell V. L. Barclay, Michael J. W. Boyle, Arthur Y. C. Chung, Tom M. Fayle, Huai En Hah, Jane L. Hardwick, Lois Kinneen, Roger L. Kitching, Sarah C. Maunsell, Jeremy A. Miller, Adam C. Sharp, Nigel E. Stork, Leona Wai, Kalsum M. Yusah, Robert M. Ewers. (2023). Tropical invertebrate community assembly processes are robust to a gradient of land use intensity. https://doi.org/10.1101/2023.01.30.526177

