## Supplementary figures and tables for "Tropical invertebrate community assembly processes are robust to a gradient of land use intensity"

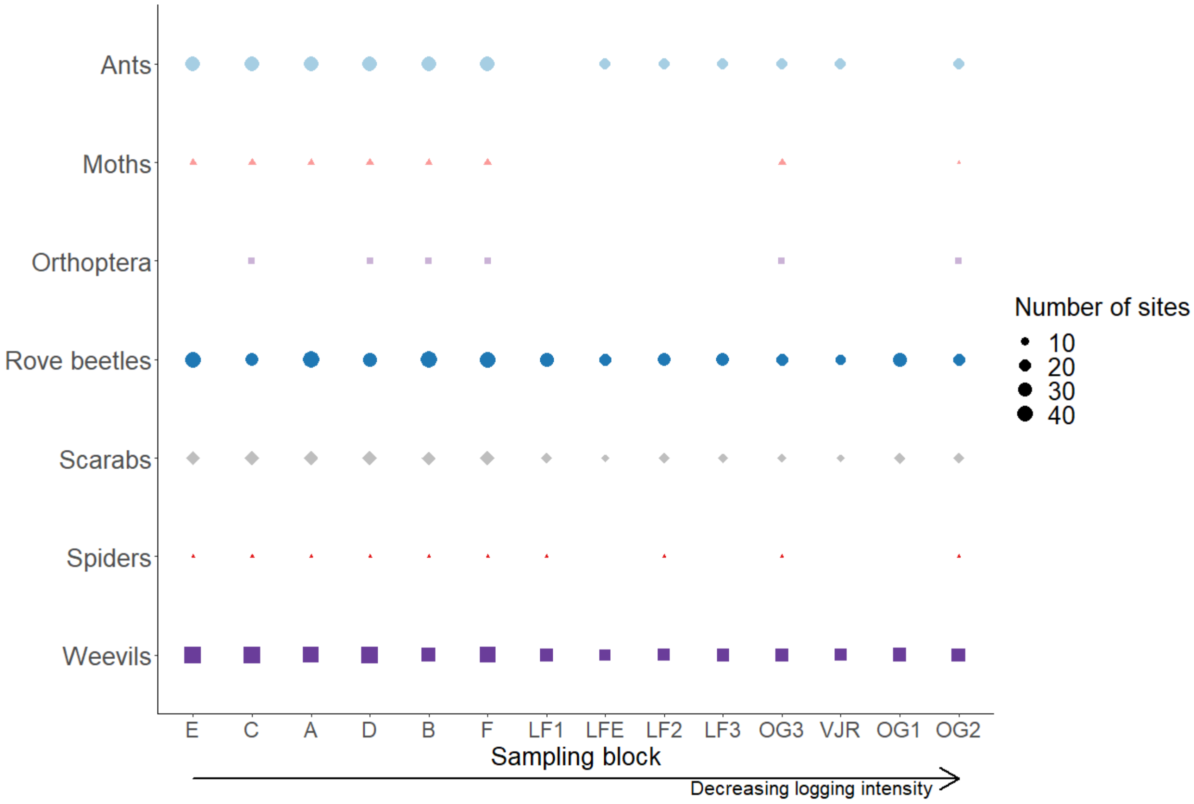


**Figure S1.** Sample sizes across the different taxa included in the study. Each point shows that the taxa were sampled in that block. The size of the points corresponds to the number of sites sampled in that block, where the minimum number of sites is 4 for spiders, and the maximum number of sites is 43 at block E for weevils. Blocks named A-F represent logged forest fragments. LF is logged forest, OG is old-growth forest, and VJR is virgin jungle reserve. Blocks are presented along the *x* axis in order of increasing above-ground carbon density (ACD). However, for scarabs and spiders, block C had a slightly lower average ACD than block E. Also, for weevils, block VJR had a slightly lower ACD than block OG3.

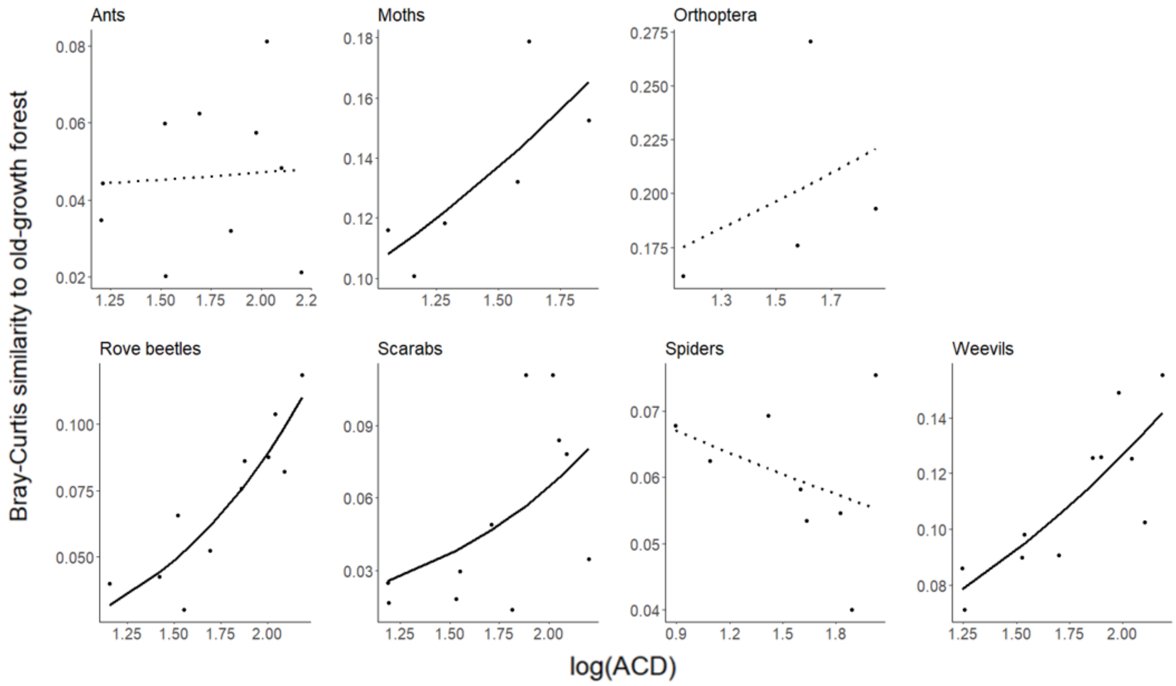


**Figure S2.** Mean Bray-Curtis community similarity to old-growth forest. Each point represents one sampling block, shown in order of increasing above-ground carbon density (ACD) on the *x* axis. The species composition of each sampling block was compared to that of old-growth forest. Slopes were calculated using beta regression. Significant slopes are shown with solid lines and non-significant slopes with dotted lines.

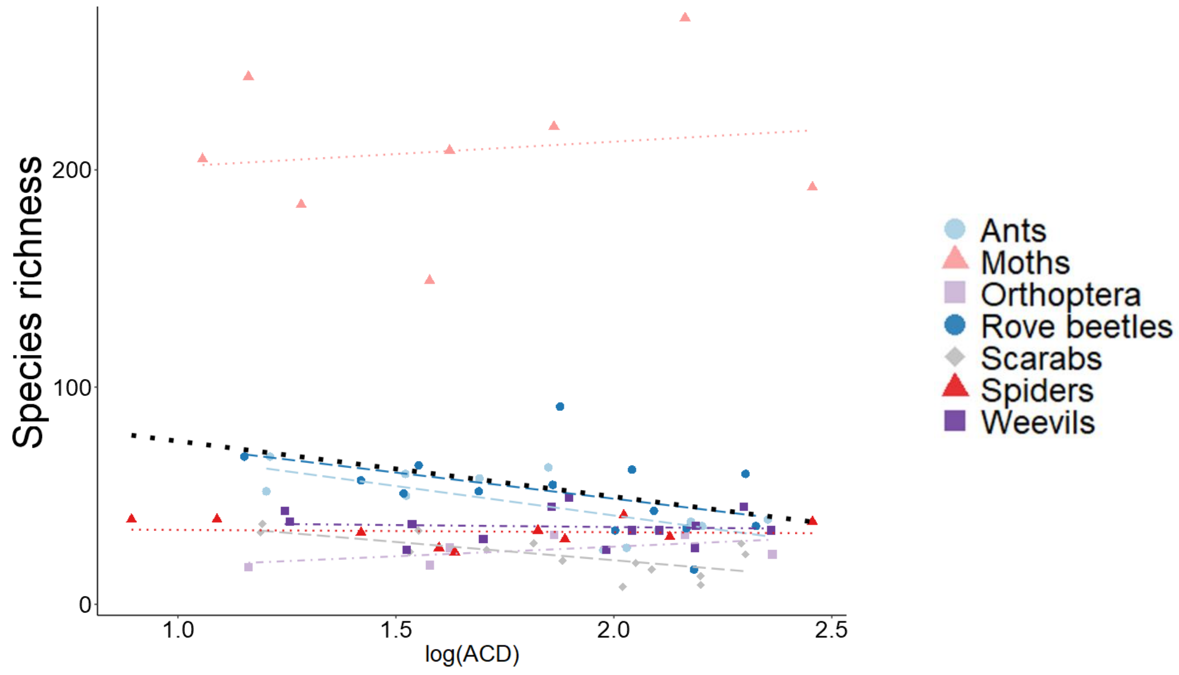


**Figure S3.** Average species richness in each sampling block for each taxon. Each point represents one sampling block, shown in order of increasing above-ground carbon density (ACD) on the x axis.

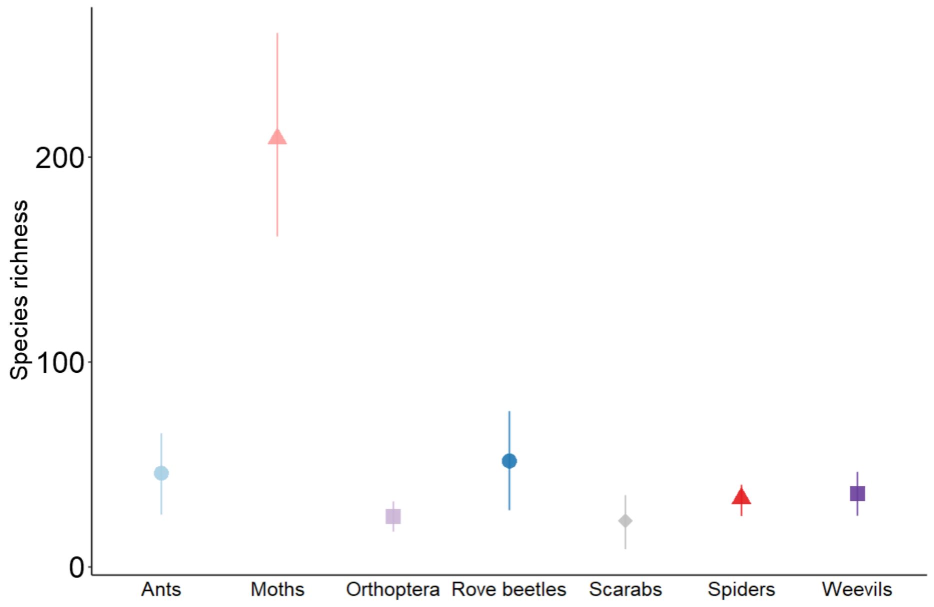


**Figure S4.** Species richness averaged across all blocks for each taxon.

**Table S1.** Summary of community composition data.

| Group of taxa | Date of data collection | Sampling method | Number of sampling blocks | Number of sites | Number of species | Number of individuals | DOI |
|---|---|---|---|---|---|---|---|
| Ants | 2011-2012 | Bait cards | 14 | 296 | 244 | 18,512 | 10.5281/zenodo.3876227 |
| Moths | 2014 | UV light traps | 8 | 46 | 643 | 6,447 | 10.5281/zenodo.4247169 |
| Orthoptera | 2015 | Sweep netting | 6 | 36 | 64 | 747 | 10.5281/zenodo.4275386 |
| Rove beetles | 2011-2013 | Combination pitfall-malaise traps | 14 | 384 | 252 | 2,656 | 10.5281/zenodo.1323504 |
| Scarabs | 2011-2013 | Combination pitfall-malaise traps | 14 | 299 | 112 | 673 | 10.5281/zenodo.1323504 |
| Spiders | 2015 | Beating plant foliage | 10 | 40 | 176 | 525 | 10.5281/zenodo.4139685 |
| Weevils | 2011-2013 | Combination pitfall-malaise traps | 14 | 407 | 154 | 2,734 | 10.5281/zenodo.1323504 |
| | | | | Total | 1,645 | 32,294 | |

**Table S2.** Number of (morpho)species in each phylogenetic bin used for the iCAMP analysis. Phylogenetic binning was done using the *iCAMP* package (Ning et al., 2020) with 9 species set as the minimum bin size.

| | **Number of bins** | **Total number of species** | **Numbers of species per bin** |
|---|---|---|---|
| **Ants** | 6 | 244 | 162, 14, 9, 14, 34, 11 |
| **Moths** | 5 | 643 | 400, 34, 19, 78, 14 |
| **Orthoptera** | 2 | 64 | 45, 15 |
| **Rove beetles** | 7 | 252 | 177, 19, 9, 10, 9, 9, 19 |
| **Scarabs** | 5 | 112 | 20, 52, 19, 9, 12 |
| **Spiders** | 5 | 176 | 50, 41, 57, 17, 11 |
| **Weevils** | 2 | 154 | 145, 9 |

**Table S3.** Overall unweighted average stochasticity metrics for the entire metacommunity (all sites pooled together, irrespective of ACD) for each group of taxa. NST is normalised stochasticity ratio and MST is modified stochasticity ratio (Ning et al., 2019). NTP is the abundance-weighted percentage of taxa whose observed occurrence frequencies are within one confidence intervals of that which would be expected by Sloan's neutral model (Sloan *et al.* 2006, 2007; Burns *et al.* 2016). Higher values indicate a greater relative importance of stochasticity.

| | **NST** | **MST** | **NTP** |
|---|---|---|---|
| **Ants** | 0.67 | 0.21 | 67 % |
| **Moths** | 0.66 | 0.65 | 61 % |
| **Orthoptera** | 0.62 | 0.54 | 59 % |
| **Rove beetles** | 0.38 | 0.13 | 77 % |
| **Scarabs** | 0.16 | 0.02 | NA |
| **Spiders** | 0.58 | 0.47 | 70 % |
| **Weevils** | 0.54 | 0.26 | 65 % |

**Table S4.** Overall relative importance of community assembly processes calculated by iCAMP.

| Process | Group of taxa | Mean relative importance of process (%) | Lower 95% quantile (LCI) of mean (%) | Upper 95% quantile (UCI) of mean (%) | Weight = 1/ (UCI – LCI) |
|---|---|---|---|---|---|
| **Heterogeneous selection** | Ants | 4.0 | 2.0 | 6.0 | 25.00 |
| | Moths | 14.9 | 6.7 | 17.4 | 9.32 |
| | Orthoptera | 18.6 | 19.2 | 18.0 | -83.98 |
| | Rove beetles | 0.1 | 0.0 | 0.2 | 525.31 |
| | Scarabs | 0.0 | 0.0 | 0.0 | 0.00 |
| | Spiders | 0.0 | 0.0 | 0.0 | 0.00 |
| | Weevils | 0.6 | 0.0 | 4.1 | 24.36 |
| | **Weighted summary mean for all taxa = -2.6 %, CI = [0.1, 18.6]** | | | | |
| **Homogeneous selection** | Ants | 1.0 | 1.0 | 2.0 | 100.00 |
| | Moths | 0.2 | 0.1 | 0.3 | 342.97 |
| | Orthoptera | 0.1 | 0.0 | 0.2 | 512.44 |
| | Rove beetles | 5.9 | 3.8 | 12.7 | 11.19 |
| | Scarabs | 3.0 | 0.7 | 4.3 | 28.01 |
| | Spiders | 4.7 | 0.2 | 8.0 | 12.76 |
| | Weevils | 0.4 | 0.0 | 1.1 | 97.40 |
| | **Weighted summary mean for all taxa = 0.4 %, CI = [0.4, 5.7]** | | | | |
| **Dispersal limitation** | Ants | 95.0 | 93.0 | 97.0 | 25.00 |
| | Moths | 13.4 | 7.2 | 19.9 | 7.87 |
| | Orthoptera | 3.8 | 1.4 | 7.1 | 17.71 |
| | Rove beetles | 93.9 | 88.0 | 97.0 | 11.09 |
| | Scarabs | 96.9 | 95.9 | 99.3 | 29.77 |
| | Spiders | 5.9 | 2.5 | 8.2 | 17.54 |
| | Weevils | 96.2 | 90.2 | 98.1 | 12.58 |
| | **Weighted summary mean for all taxa = 64.1 %, CI = [25.5, 96.9]** | | | | |
| **Homogenising dispersal** | Ants | 0.0 | 0.0 | 0.0 | 0.00 |
| | Moths | 4.4 | 0.4 | 9.2 | 11.28 |
| | Orthoptera | 2.4 | 0.5 | 5.0 | 22.32 |
| | Rove beetles | 0.0 | 0.0 | 0.0 | 0.00 |
| | Scarabs | 0.0 | 0.0 | 0.0 | 0.00 |
| | Spiders | 0.5 | 0.0 | 1.2 | 83.99 |

| | | | | | |
|---|---|---|---|---|---|
| | Weevils | 0.0 | 0.0 | 0.0 | 0.00 |
| | **Weighted summary mean for all taxa = 1.2 %, CIs = [0.0, 2.1]** | | | | |
| **Drift** | Ants | 0.0 | 0.0 | 0.0 | 0.00 |
| | Moths | 67.2 | 54.2 | 77.8 | 4.23 |
| | Orthoptera | 75.1 | 55.3 | 87.0 | 3.16 |
| | Rove beetles | 0.2 | 0.0 | 0.6 | 156.58 |
| | Scarabs | 0.0 | 0.0 | 0.1 | 1838.90 |
| | Spiders | 88.9 | 77.5 | 95.7 | 5.50 |
| | Weevils | 2.7 | 0.8 | 7.6 | 14.72 |
| | **Weighted summary mean for all taxa = 0.5 %, CIs = [0.0, 75.1]** | | | | |

**Table S5.** Slopes of the regressions where above-ground carbon density, taxa and their interaction are predictor variables and the relative importances of each community assembly process are the response variables.

| Process | Group of taxa | Mean slope | Lower 95% quantile (LCI) of slope | Upper 95% quantile (UCI) of slope | Weight = 1/ (UCI – LCI) |
|---|---|---|---|---|---|
| **Heterogeneous selection** | Ants | 0.33 | -0.27 | 0.93 | 0.83 |
| | Moths | 2.45 | 1.79 | 3.11 | 0.76 |
| | Orthoptera | 1.73 | -0.19 | 3.65 | 0.26 |
| | Rove beetles | 0.16 | -1.94 | 2.26 | 0.24 |
| | Scarabs | 0.00 | -2.06 | 2.06 | 0.24 |
| | Spiders | -0.35 | -2.50 | 1.81 | 0.23 |
| | Weevils | -0.51 | -2.39 | 1.37 | 0.27 |
| | **Weighted summary mean slope for all taxa = 0.85, CI = [-0.32, 2.01]** | | | | |
| **Homogeneous selection** | Ants | -0.24 | 0.79 | -1.27 | -0.49 |
| | Moths | -1.25 | -2.56 | 0.07 | 0.38 |
| | Orthoptera | -0.50 | -3.93 | 2.93 | 0.15 |
| | Rove beetles | 1.12 | -2.16 | 4.40 | 0.15 |
| | Scarabs | -0.13 | -3.37 | 3.11 | 0.15 |
| | Spiders | -1.20 | -4.39 | 1.99 | 0.16 |
| | Weevils | 0.71 | -2.65 | 4.07 | 0.15 |
| | **Weighted summary mean slope for all taxa = -0.55 CI = [-2.98, 1.87]** | | | | |
| **Dispersal limitation** | Ants | -0.33 | -0.93 | 0.27 | 0.83 |
| | Moths | -2.45 | -3.11 | -1.79 | 0.76 |
| | Orthoptera | -1.73 | -3.65 | 0.19 | 0.26 |

| | | | | | |
|---|---|---|---|---|---|
| | Rove beetles | -0.16 | -2.26 | 1.94 | 0.24 |
| | Scarabs | 0.00 | -2.06 | 2.06 | 0.24 |
| | Spiders | 0.35 | -1.81 | 2.50 | 0.23 |
| | Weevils | 0.51 | -1.37 | 2.39 | 0.27 |
| | **Weighted summary mean slope for all groups of taxa = -0.85, CI = [-2.01, 0.32]** | | | | |
| **Homogenising dispersal** | Ants | -0.31 | -1.67 | -1.27 | 2.50 |
| | Moths | -2.49 | -3.34 | -1.64 | 0.59 |
| | Orthoptera | -1.83 | -4.09 | 0.42 | 0.22 |
| | Rove beetles | -0.08 | -2.93 | 2.78 | 0.18 |
| | Scarabs | 0.00 | -2.78 | 2.78 | 0.18 |
| | Spiders | -0.34 | -2.95 | 2.27 | 0.19 |
| | Weevils | 0.50 | -2.18 | 3.17 | 0.19 |
| | **Weighted summary mean slope for all groups of taxa = -0.65, CIs = [-1.63, 0.32]** | | | | |
| **Drift** | Ants | -0.30 | -1.66 | 1.06 | 0.37 |
| | Moths | -1.53 | -1.97 | -1.08 | 1.13 |
| | Orthoptera | -0.25 | -1.37 | 0.87 | 0.45 |
| | Rove beetles | 0.06 | -2.26 | 2.38 | 0.22 |
| | Scarabs | 0.01 | -2.21 | 2.24 | 0.22 |
| | Spiders | 0.22 | -1.51 | 1.95 | 0.29 |
| | Weevils | 0.26 | -1.55 | 2.06 | 0.28 |
| | **Weighted summary mean for all groups of taxa = -0.61, CIs = [-1.75, 0.53]** | | | | |

**Text S1**: Sensitivity analysis for spiders, Orthoptera and moths.

To test for an under-sampling effect in the Orthoptera and spiders data, we grouped together sites that were in the same sampling block and re-analysed the community assembly. Each row in the site x species matrix represented an entire sampling block. Each sampling block includes six sites for the Orthoptera data, four sites for the spiders data and six sites for the moths data (except one block, OG2, in the moths data that had four sites).

Ecological drift was still the dominant community assembly process for spiders, Orthoptera and moths. Drift underpinned 72 % (95 % quantiles = 0 – 100) of Orthoptera community assembly when all sites within the same sampling block were grouped together, compared to 74 % (95 % quantiles = 60 – 82) when each site was a separate row in the site x species matrix. Grouping sites by sampling block for Orthoptera increased the importance of heterogeneous selection from 19 % (95 % quantiles = 14 – 26) to 29 % (95 % quantiles = 0 – 100). When sites in the spiders data were grouped by sampling block, the relative importance of drift reduced from 92 % (95 % quantiles = 86– 94) to 74 % (95 % quantiles = 41 – 100), and the relative importance of homogeneous selection increased from 3 % (95 % quantiles = 1 – 3) to 15 % (95 % quantiles = 0 – 60). Grouping sites within the same block increased the relative importance of drift in moth communities from 67 % (95 % quantiles = 62 – 75) to 71 % (95 % quantiles = 4.8 – 100).

Phylogenetic trees for all groups of taxa included in this study. Bins show the phylogenetic groupings used for analysing the relative importance of community assembly processes with iCAMP. Taxonomic identity is used as a proxy for phylogenetic relatedness.

### Ants

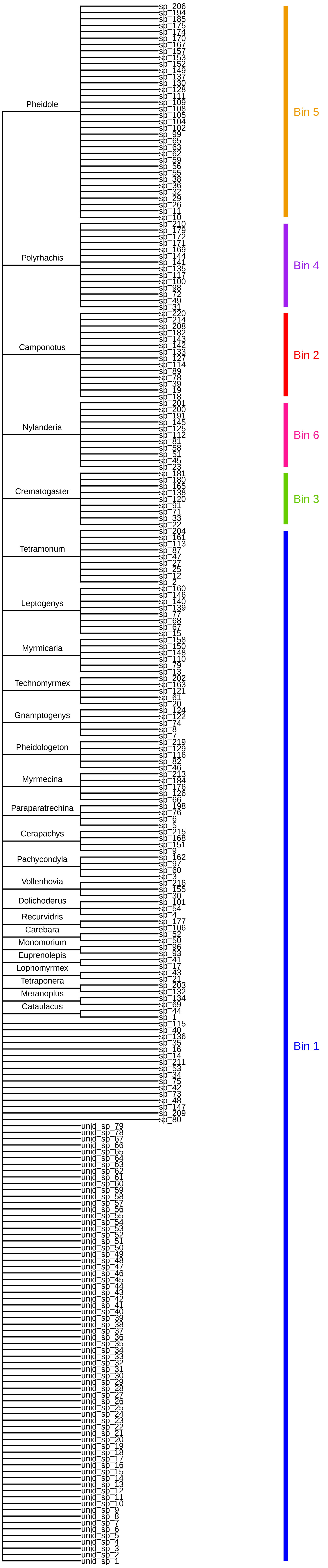

Moths

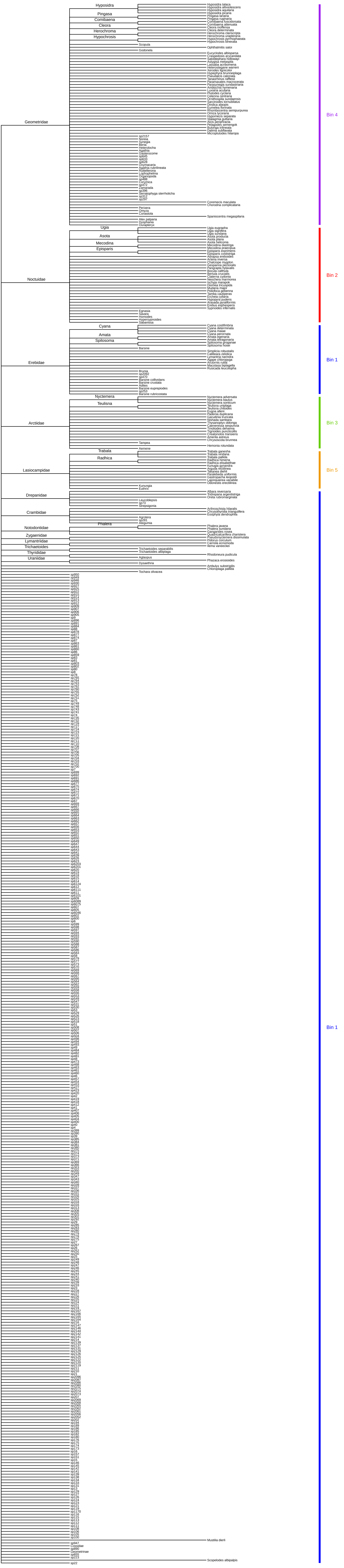

Geometridae
Hyposidra
Pingasa
Comibaena
Cleora
Herochroma
Hypochrosis
Noctuidae
Ugia
Asota
Mecodina
Episparis
Erebidae
Cyana
Amata
Spilosoma
Arctiidae
Nyctemera
Teulisna
Lasiocampidae
Trabala
Radhica
Drepanidae
Crambidae
Notodontidae
Phalera
Zygaenidae
Lymantriidae
Trichaetoides
Thyrididae
Uraniidae
Bin 4
Bin 2
Bin 1
Bin 3
Bin 5
Bin 1

Orthoptera

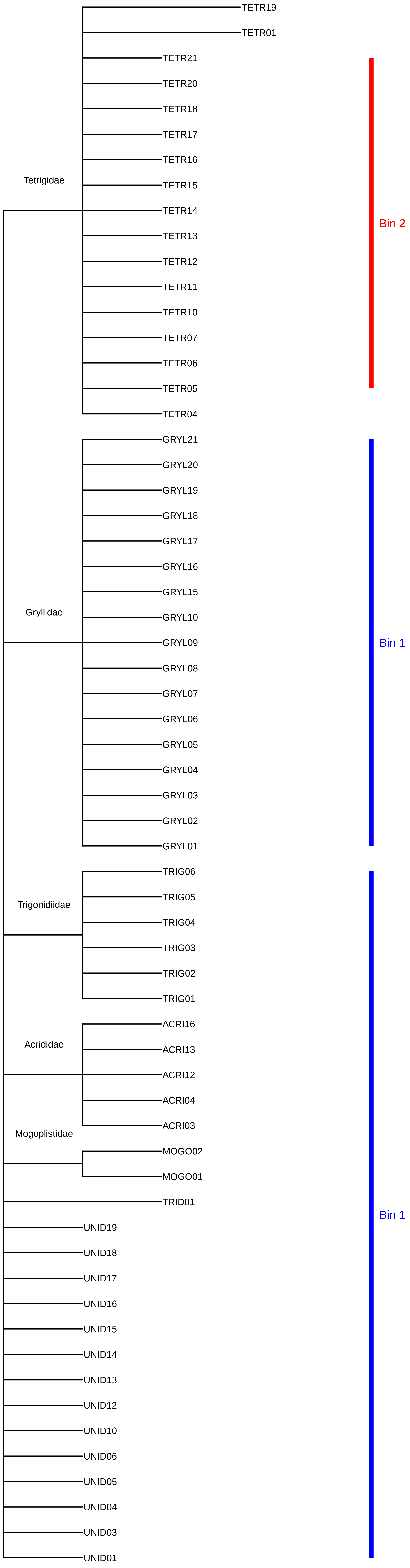

Tetrigidae
TETR19
TETR01
TETR21
TETR20
TETR18
TETR17
TETR16
TETR15
TETR14
TETR13
TETR12
TETR11
TETR10
TETR07
TETR06
TETR05
TETR04
Bin 2
Gryllidae
GRYL21
GRYL20
GRYL19
GRYL18
GRYL17
GRYL16
GRYL15
GRYL10
GRYL09
GRYL08
GRYL07
GRYL06
GRYL05
GRYL04
GRYL03
GRYL02
GRYL01
Bin 1
Trigonidiidae
TRIG06
TRIG05
TRIG04
TRIG03
TRIG02
TRIG01
Acrididae
ACRI16
ACRI13
ACRI12
ACRI04
ACRI03
Mogoplistidae
MOGO02
MOGO01
TRID01
UNID19
UNID18
UNID17
UNID16
UNID15
UNID14
UNID13
UNID12
UNID10
UNID06
UNID05
UNID04
UNID03
UNID01
Bin 1

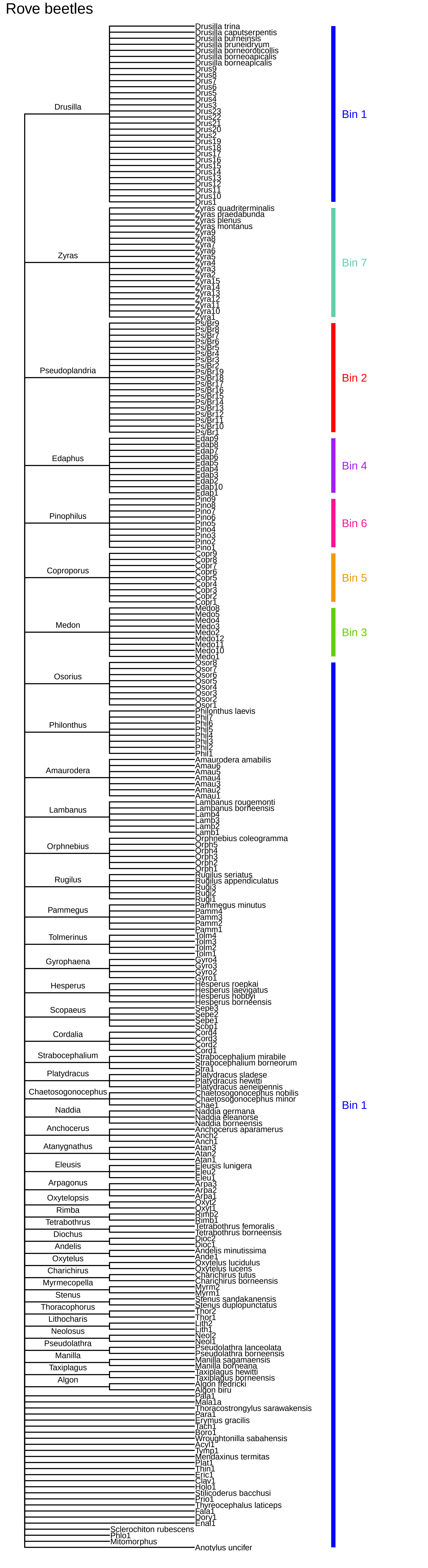
Rove beetles
Drusilla
Drusilla trina
Drusilla caputserpentis
Drusilla burneinsis
Drusilla bruneidryum
Drusilla borneoroticollis
Drusilla borneoapicalis
Drusilla borneapicalis
Drus9
Drus8
Drus7
Drus6
Drus5
Drus4
Drus3
Drus23
Drus22
Drus21
Drus20
Drus2
Drus19
Drus18
Drus17
Drus16
Drus15
Drus14
Drus13
Drus12
Drus11
Drus10
Drus1
Bin 1
Zyras
Zyras quadriterminalis
Zyras praedabunda
Zyras plenus
Zyras montanus
Zyra9
Zyra8
Zyra7
Zyra6
Zyra5
Zyra4
Zyra3
Zyra2
Zyra15
Zyra14
Zyra13
Zyra12
Zyra11
Zyra10
Zyra1
Bin 7
Pseudoplandria
Ps/Br9
Ps/Br8
Ps/Br7
Ps/Br6
Ps/Br5
Ps/Br4
Ps/Br3
Ps/Br2
Ps/Br19
Ps/Br18
Ps/Br17
Ps/Br16
Ps/Br15
Ps/Br14
Ps/Br13
Ps/Br12
Ps/Br11
Ps/Br10
Ps/Br1
Bin 2
Edaphus
Edap9
Edap8
Edap7
Edap6
Edap5
Edap4
Edap3
Edap2
Edap10
Edap1
Bin 4
Pinophilus
Pino9
Pino8
Pino7
Pino6
Pino5
Pino4
Pino3
Pino2
Pino1
Bin 6
Coproporus
Copr9
Copr8
Copr7
Copr6
Copr5
Copr4
Copr3
Copr2
Copr1
Bin 5
Medon
Medo8
Medo5
Medo4
Medo3
Medo2
Medo12
Medo11
Medo10
Medo1
Bin 3
Osorius
Osor8
Osor7
Osor6
Osor5
Osor4
Osor3
Osor2
Osor1
Philonthus
Philonthus laevis
Phil7
Phil6
Phil5
Phil4
Phil3
Phil2
Phil1
Amaurodera
Amaurodera amabilis
Amau6
Amau5
Amau4
Amau3
Amau2
Amau1
Lambanus
Lambanus rougemonti
Lambanus borneensis
Lamb4
Lamb3
Lamb2
Lamb1
Orphnebius
Orphnebius coleogramma
Orph5
Orph4
Orph3
Orph2
Orph1
Rugilus
Rugilus seriatus
Rugilus appendiculatus
Rugi3
Rugi2
Rugi1
Pammegus
Pammegus minutus
Pamm4
Pamm3
Pamm2
Pamm1
Tolmerinus
Tolm4
Tolm3
Tolm2
Tolm1
Gyrophaena
Gyro4
Gyro3
Gyro2
Gyro1
Hesperus
Hesperus roepkai
Hesperus laevigatus
Hesperus hobbyi
Hesperus borneensis
Scopaeus
Sepe3
Sepe2
Sepe1
Scop1
Cordalia
Cord4
Cord3
Cord2
Cord1
Strabocephalium
Strabocephalium mirabile
Strabocephalium borneorum
Stra1
Platydracus
Platydracus sladese
Platydracus hewitti
Platydracus aeneipennis
Chaetosogonocephus
Chaetosogonocephus nobilis
Chaetosogonocephus minor
Chae1
Naddia
Naddia germana
Naddia eleanorse
Naddia borneensis
Anchocerus
Anchocerus aparamerus
Anch2
Anch1
Atanygnathus
Atan3
Atan2
Atan1
Eleusis
Eleusis lunigera
Eleu2
Eleu1
Arpagonus
Arpa3
Arpa2
Arpa1
Oxytelopsis
Oxyt2
Oxyt1
Rimba
Rimb2
Rimb1
Tetrabothrus
Tetrabothrus femoralis
Tetrabothrus borneensis
Diochus
Dioc2
Dioc1
Andelis
Andelis minutissima
Ande1
Oxytelus
Oxytelus lucidulus
Oxytelus lucens
Charichirus
Charichirus tutus
Charichirus borneensis
Myrmecopella
Myrm2
Myrm1
Stenus
Stenus sandakanensis
Stenus duplopunctatus
Thoracophorus
Thor2
Thor1
Lithocharis
Lith2
Lith1
Neolosus
Neol2
Neol1
Pseudolathra
Pseudolathra lanceolata
Pseudolathra borneensis
Manilla
Manilla sagamaensis
Manilla borneana
Taxiplagus
Taxiplagus hewitti
Taxiplagus borneensis
Algon
Algon fredricki
Algon biru
Pala1
Mala1a
Thoracostrongylus sarawakensis
Para1
Erymus gracilis
Tach1
Boro1
Wroughtonilla sabahensis
Acyl1
Tymp1
Mendaxinus termitas
Plat1
Thin1
Eric1
Clav1
Holo1
Stilicoderus bacchui
Prio1
Thyreocephalus laticeps
Fala1
Dory1
Enal1
Sclerochiton rubescens
Phlo1
Mitomorphus
Anotylus uncifer
Bin 1

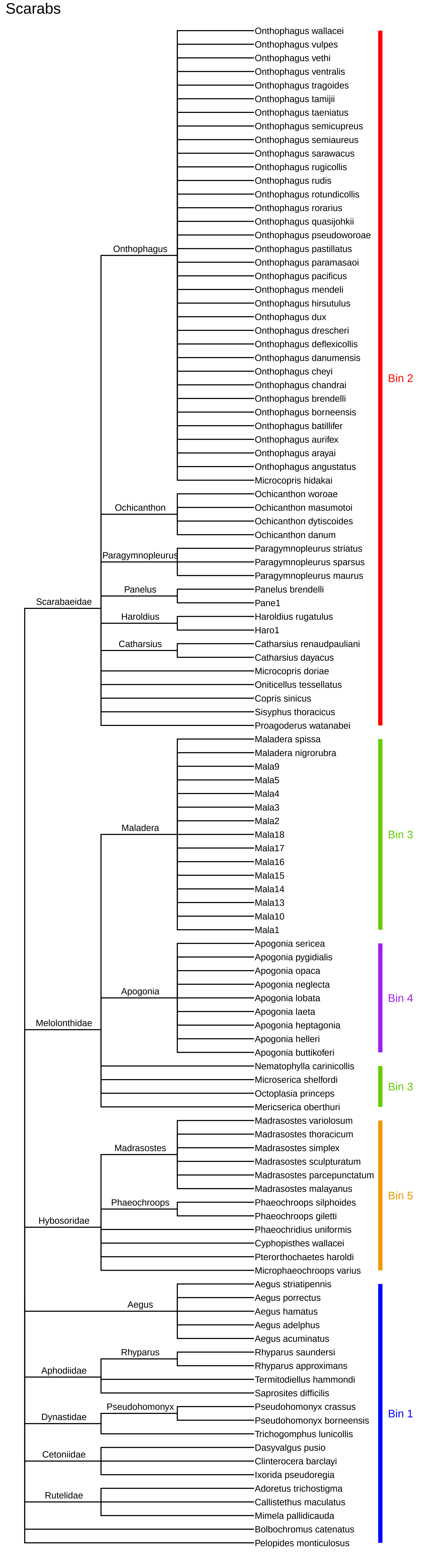
Scarabs
Scarabaeidae
Onthophagus
Onthophagus wallacei
Onthophagus vulpes
Onthophagus vethi
Onthophagus ventralis
Onthophagus tragoides
Onthophagus tamijii
Onthophagus taeniatus
Onthophagus semicupreus
Onthophagus semiaureus
Onthophagus sarawacus
Onthophagus rugicollis
Onthophagus rudis
Onthophagus rotundicollis
Onthophagus rorarius
Onthophagus quasijohkii
Onthophagus pseudoworoae
Onthophagus pastillatus
Onthophagus paramasaoi
Onthophagus pacificus
Onthophagus mendeli
Onthophagus hirsutulus
Onthophagus dux
Onthophagus drescheri
Onthophagus deflexicollis
Onthophagus danumensis
Onthophagus cheyi
Onthophagus chandrai
Onthophagus brendelli
Onthophagus borneensis
Onthophagus batillifer
Onthophagus aurifex
Onthophagus arayai
Onthophagus angustatus
Microcopris hidakai
Ochicanthon
Ochicanthon woroae
Ochicanthon masumotoi
Ochicanthon dytiscoides
Ochicanthon danum
Paragymnopleurus
Paragymnopleurus striatus
Paragymnopleurus sparsus
Paragymnopleurus maurus
Panelus
Panelus brendelli
Pane1
Haroldius
Haroldius rugatulus
Haro1
Catharsius
Catharsius renaudpauliani
Catharsius dayacus
Microcopris doriae
Oniticellus tessellatus
Copris sinicus
Sisyphus thoracicus
Proagoderus watanabei
Bin 2
Melolonthidae
Maladera
Maladera spissa
Maladera nigrorubra
Mala9
Mala5
Mala4
Mala3
Mala2
Mala18
Mala17
Mala16
Mala15
Mala14
Mala13
Mala10
Mala1
Bin 3
Apogonia
Apogonia sericea
Apogonia pygidialis
Apogonia opaca
Apogonia neglecta
Apogonia lobata
Apogonia laeta
Apogonia heptagonia
Apogonia helleri
Apogonia buttikoferi
Bin 4
Nematophylla carinicollis
Microserica shelfordi
Octoplasia princeps
Mericserica oberthuri
Bin 3
Hybosoridae
Madrasostes
Madrasostes variolosum
Madrasostes thoracicum
Madrasostes simplex
Madrasostes sculpturatum
Madrasostes parcepunctatum
Madrasostes malayanus
Phaeochroops
Phaeochroops silphoides
Phaeochroops giletti
Phaeochridius uniformis
Cyphopisthes wallacei
Pterorthochaetes haroldi
Microphaeochroops varius
Bin 5
Aegus
Aegus striatipennis
Aegus porrectus
Aegus hamatus
Aegus adelphus
Aegus acuminatus
Aphodiidae
Rhyparus
Rhyparus saundersi
Rhyparus approximans
Termitodiellus hammondi
Saprosites difficilis
Dynastidae
Pseudohomonyx
Pseudohomonyx crassus
Pseudohomonyx borneensis
Trichogomphus lunicollis
Cetoniidae
Dasyvalgus pusio
Clinterocera barclayi
Ixorida pseudoregia
Rutelidae
Adoretus trichostigma
Callistethus maculatus
Mimela pallidicauda
Bolbochromus catenatus
Pelopides monticulosus
Bin 1

Spiders

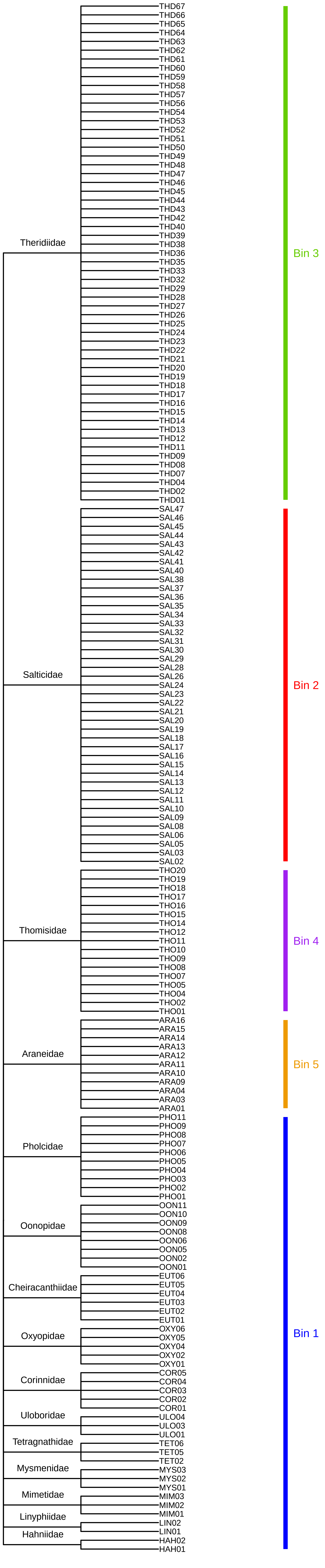
Theridiidae
Salticidae
Thomisidae
Araneidae
Pholcidae
Oonopidae
Cheiracanthiidae
Oxyopidae
Corinnidae
Uloboridae
Tetragnathidae
Mysmenidae
Mimetidae
Linyphiidae
Hahniidae
THD67
THD66
THD65
THD64
THD63
THD62
THD61
THD60
THD59
THD58
THD57
THD56
THD54
THD53
THD52
THD51
THD50
THD49
THD48
THD47
THD46
THD45
THD44
THD43
THD42
THD40
THD39
THD38
THD36
THD35
THD33
THD32
THD29
THD28
THD27
THD26
THD25
THD24
THD23
THD22
THD21
THD20
THD19
THD18
THD17
THD16
THD15
THD14
THD13
THD12
THD11
THD09
THD08
THD07
THD04
THD02
THD01
SAL47
SAL46
SAL45
SAL44
SAL43
SAL42
SAL41
SAL40
SAL38
SAL37
SAL36
SAL35
SAL34
SAL33
SAL32
SAL31
SAL30
SAL29
SAL28
SAL26
SAL24
SAL23
SAL22
SAL21
SAL20
SAL19
SAL18
SAL17
SAL16
SAL15
SAL14
SAL13
SAL12
SAL11
SAL10
SAL09
SAL08
SAL06
SAL05
SAL03
SAL02
THO20
THO19
THO18
THO17
THO16
THO15
THO14
THO12
THO11
THO10
THO09
THO08
THO07
THO05
THO04
THO02
THO01
ARA16
ARA15
ARA14
ARA13
ARA12
ARA11
ARA10
ARA09
ARA04
ARA03
ARA01
PHO11
PHO09
PHO08
PHO07
PHO06
PHO05
PHO04
PHO03
PHO02
PHO01
OON11
OON10
OON09
OON08
OON06
OON05
OON02
OON01
EUT06
EUT05
EUT04
EUT03
EUT02
EUT01
OXY06
OXY05
OXY04
OXY02
OXY01
COR05
COR04
COR03
COR02
COR01
ULO04
ULO03
ULO01
TET06
TET05
TET02
MYS03
MYS02
MYS01
MIM03
MIM02
MIM01
LIN02
LIN01
HAH02
HAH01
Bin 3
Bin 2
Bin 4
Bin 5
Bin 1

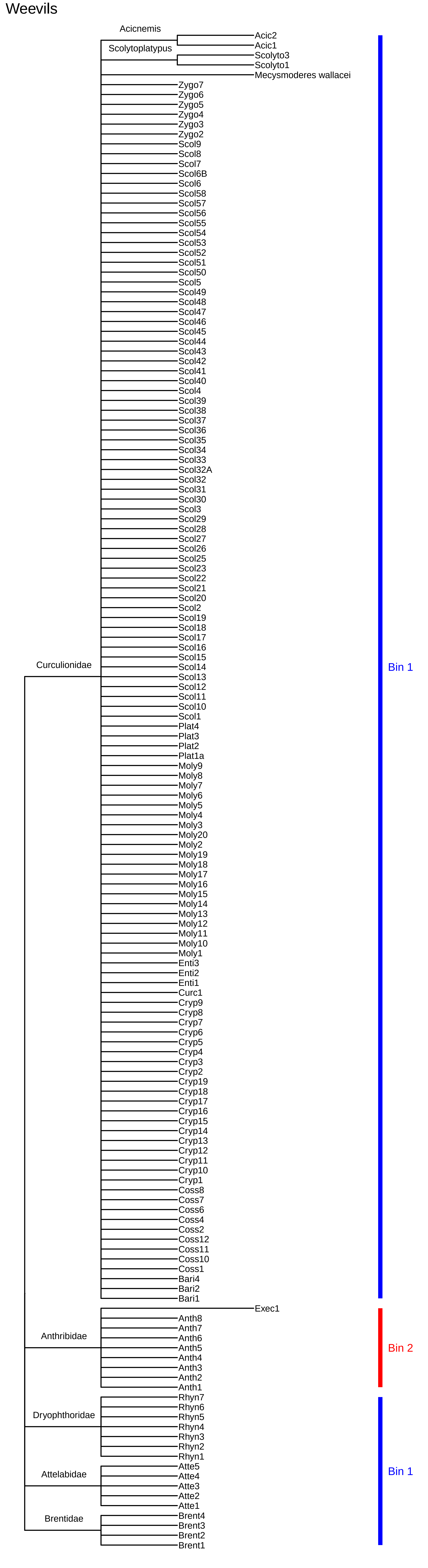
Weevils
Acicnemis
Scolytoplatypus
Acic2
Acic1
Scolyto3
Scolyto1
Mecysmoderes wallacei
Zygo7
Zygo6
Zygo5
Zygo4
Zygo3
Zygo2
Scol9
Scol8
Scol7
Scol6B
Scol6
Scol58
Scol57
Scol56
Scol55
Scol54
Scol53
Scol52
Scol51
Scol50
Scol5
Scol49
Scol48
Scol47
Scol46
Scol45
Scol44
Scol43
Scol42
Scol41
Scol40
Scol4
Scol39
Scol38
Scol37
Scol36
Scol35
Scol34
Scol33
Scol32A
Scol32
Scol31
Scol30
Scol3
Scol29
Scol28
Scol27
Scol26
Scol25
Scol23
Scol22
Scol21
Scol20
Scol2
Scol19
Scol18
Scol17
Scol16
Scol15
Scol14
Scol13
Scol12
Scol11
Scol10
Scol1
Plat4
Plat3
Plat2
Plat1a
Moly9
Moly8
Moly7
Moly6
Moly5
Moly4
Moly3
Moly20
Moly2
Moly19
Moly18
Moly17
Moly16
Moly15
Moly14
Moly13
Moly12
Moly11
Moly10
Moly1
Enti3
Enti2
Enti1
Curc1
Cryp9
Cryp8
Cryp7
Cryp6
Cryp5
Cryp4
Cryp3
Cryp2
Cryp19
Cryp18
Cryp17
Cryp16
Cryp15
Cryp14
Cryp13
Cryp12
Cryp11
Cryp10
Cryp1
Coss8
Coss7
Coss6
Coss4
Coss2
Coss12
Coss11
Coss10
Coss1
Bari4
Bari2
Bari1
Curculionidae
Bin 1
Exec1
Anth8
Anth7
Anth6
Anth5
Anth4
Anth3
Anth2
Anth1
Anthribidae
Bin 2
Rhyn7
Rhyn6
Rhyn5
Rhyn4
Rhyn3
Rhyn2
Rhyn1
Dryophthoridae
Atte5
Atte4
Atte3
Atte2
Atte1
Attelabidae
Bin 1
Brent4
Brent3
Brent2
Brent1
Brentidae
